# *Ab-initio* discovery of tumoricidal oligonucleotides in a DNA sequencing machine

**DOI:** 10.1101/630830

**Authors:** Noam Mamet, Itai Rusinek, Gil Harari, Zvi Shapira, Yaniv Amir, Erez Lavi, Adva Zamir, Noam Borovsky, Noah Joseph, Maria Motin, Dekel Saban, Ido Bachelet

**Affiliations:** Molecular Computing Cluster, Augmanity, Rehovot, Israel. Website: https://augm.com; Faculty of Life Sciences, Bar-Ilan University, Israel

## Abstract

We describe a technique for the rapid *ab-initio* discovery of target-tailored tumoricidal DNA oligonucleotides inside an Illumina sequencing chip. By sequencing oligonucleotide pools we generate a physical microfluidic map of hundreds of millions of potential oligo clusters, in which every cluster is mapped to a specific set of spatial coordinates. Tumor cells, pre-loaded with a fluorogenic reporter of apoptosis, are then injected into the chip and monitored over time. Apoptotic tumor cells are identified and analyzed across the entire map, automatically revealing the coordinates of oligos that induced this effect. We demonstrate this method by identifying, within just a few hours, new oligos capable of directly and selectively inducing apoptosis in primary human tumor cells. Such a major capability could lead to a new paradigm of personalized cancer therapy.

## Introduction

Drug discovery - the ongoing search for better molecular and cellular tools to treat disease - is a major process propelling medicine forward. The products of this search have been constantly improving in the past century towards safer and more effective therapeutics. In particular, the relatively recent introduction of personalized therapeutic technologies arguably represents a significant conceptual breakthrough^1,2^. This premise is of particular importance in cancer, which exhibits high variability and patient heterogeneity^3^, with tumors constantly developing resistance to drugs, requiring dynamic treatment adaptation. It is nearly impossible for a single drug or protocol to be both effective and safe across many patients^4^, which strongly highlights the necessity of personalized therapeutic strategies.

Interestingly, the term “personalized medicine” may be interpreted into several different meanings. For instance, it is often used in the context of matching a specific drug to a patient based on patient-specific information, rather than by a generic protocol. The decision to match could incorporate genetic information, observed response to a panel of drugs, etc., but in all cases the drug is selected from the set of existing ones. A completely different meaning of the term relates to procedures implemented on patient-derived material, as is the case in chimeric antigen receptor (CAR)-T cells^5^ or dendritic cell vaccines^6^. Undoubtedly, all these strategies interface with the patient in a profoundly better and smarter way than conventional generic approaches do; however, they are still confined to a single procedure or a narrow set of drugs, and could thus still be hypothetically expanded on towards entirely new and improved types of drug discovery, in which drugs are tailored *de-novo*, per case, and on-demand. Although this vision remains to be built and evaluated, it already raises fascinating questions regarding the potential candidate substrate that could support such drug discovery strategies. In this study, our aim was to evaluate high-throughput DNA sequencing as an engine to drive such a technology.

High-throughput DNA sequencing offers a unique setting to test en mass diverse functions of oligonucleotides, which utilizes the mechanics of the sequencing strategy being used^20^. For example, the Illumina technology sequences DNA by using sample oligonucleotides as templates for de-novo DNA synthesis^21^, which is performed on billions of unique sample strands in parallel. This process generates a 2D map of oligonucleotide clusters, which associates cluster sequences to physical coordinates on the microfluidic sequencing chip. This feature is essentially identical in all Illumina sequencers, and several machines, such as the Genome Analyzer (GA)IIx, MiSeq, and NextSeq, have been used, following modification of recipes and components within the chemistry and hardware of the machine, to investigate various functionalities of DNA, RNA, and even peptides. Recent examples include transcription factor binding to dsDNA^22^, RNA-protein interactions^23–25^, CRISPR-Cas DNA recognition^26,27^, antibody-peptide interactions^28,29^, and a massive simulation of a mathematical game^30^.

DNA oligonucleotides, which are the natural substrate of high-throughput DNA sequencing, are at the core of various intriguing technologies with therapeutic potential^9^. Oligonucleotide therapeutics can be divided into three distinct groups of therapeutic (or potentially therapeutic) molecules, depending on their mechanism of action:

1. Genetic: molecules operating at the message level to interfere with gene expression, e.g. RNAi, mRNA, antisense, and splice-switching oligonucleotides
2. Immunostimulatory pattern: molecules containing pathogen-associated molecular patterns, such as dsRNA or unmethylated CpG, which stimulate innate immune responses via Toll-like receptors 3 and 9, respectively
3. 3D structure: molecules with functions encoded by their 3D structure, e.g. aptamers and ribozymes

For purposes of this report we shall discuss specifically aptamers. These are nucleic acid oligonucleotides evolved and selected artificially to bind to molecular and cellular targets^7,8^. Within this diverse class, aptamers are interesting for several reasons. Despite having been described nearly three decades ago, and exhibiting significant advantages over antibodies - their protein-based functional equivalents - aptamers are still rare within the pharmaceutical mainstream, with only one aptamer approved for therapy^10^ (although additional ones are currently in clinical trials). Aptamers are ~10 fold smaller, in molecular terms, than antibodies; their synthesis is chemically-defined, precise, cell-free, and flexibly scalable; they are highly stable both thermally and on the shelf, easily modified for programmable pharmacokinetics, and they are relatively safe and non-immunogenic as reported from clinical trials^11–13^. But most pertinent to the hypothesis proposed above, the *de-novo* discovery of aptamers could be orders of magnitude faster than that of any other therapeutic molecule^14–16^. Moreover, their successful discovery might not require mechanistic understanding, as in the case of the cell-SELEX process, where extremely selective aptamers - with potential therapeutic and/or diagnostic value - have been discovered given only target cells as input for their selection process^17–19^. Remarkably, this case highlights effect and selectivity - arguably the two critical pharmacological determinants - as the sole drivers of aptamer selection, rather than mechanism (e.g. a specific protein or receptor on the cell surface). This unique potential of aptamers, as therapeutics that could be discovered both rapidly and unconstrained to a mechanism, stimulated us to search for new platforms for their discovery.

High-throughput DNA sequencing offers a unique setting to test *en mass* diverse functions of oligonucleotides, which utilizes the mechanics of the sequencing strategy being used^20^. For example, the Illumina technology sequences DNA by using sample oligonucleotides as templates for *Ab-initio* DNA synthesis^21^, which is performed on billions of unique sample strands in parallel. This process generates a 2D map of oligonucleotide clusters, which associates cluster sequences to physical coordinates on the microfluidic sequencing chip. This feature is essentially identical in all Illumina sequencers, and several machines, such as the Genome Analyzer (GA)IIx, MiSeq, and NextSeq, have been used, following modification of recipes and components within the chemistry and hardware of the machine, to investigate various functionalities of DNA, RNA, and even peptides. Recent examples include transcription factor binding to dsDNA^22^, RNA-protein interactions^23–25^, CRISPR-Cas DNA recognition^26,27^, antibody-peptide interactions^28,29^, and a massive simulation of a mathematical game^30^.

In this study, we utilized Illumina sequencing for the rapid discovery of aptamer-like oligonucleotides, that specifically target human cells, but extend beyond binding into cellular effects, using the apoptosis of tumor cells as a test case. We recently described the discovery of tumoricidal oligos by effect-direct *in-vitro* evolution^31^. Here we demonstrate a variation on this technique, that utilizes the physical chips produced by Illumina sequencing in order to identify tumoricidal oligos specific to patient-derived cells, with extremely promising results.

## Results

A schematic of the workflow is presented in Fig. 1A. The first step in the platform is the sequencing of a library or pool of oligonucleotides. These could come from various sources, including a standard cell-SELEX process against a specific cell type, and from one or more rounds; alternatively, they could be rationally designed based on known aptamers (and variants thereof) to the target cells; both sources have been used in our laboratory. We sequenced this library on an Illumina NextSeq 500 instrument. In order to be able to carry out procedures on the chip outside the sequencing machine, we custom designed a mechanical adapter (Fig. 1B)that enables both the incorporation of microfluidic pumps and optical scanning in a Nikon Eclipse Ti2-E microscope. Interestingly, DNA clusters synthesized in the chip during sequencing were readily detectable by phase contrast microscopy (Fig. 1C), most likely due to local variations in the local refraction index due to high DNA density. We introduced a set of fluorescently-labeled hybridization probes, complementary to control sequences, as anchors in the coordinate system of the microscope, to derive the correct transformations to the NextSeq coordinate system reported alongside the sequences in the FASTQ headers (Fig. 1D). In order to fine-tune this transformation and obtain sufficient accuracy, we sampled images of clusters captured in phase contrast and registered them to synthetic images of the NextSeq cluster locations. Registration was done by aggregation of several image registration packages^32,33^ and the results were used to parametrize an affine transformation of each microscope field of view in the microscope stage continuous coordinate system, to the corresponding NextSeq tiles (Fig. 1E). The transformation is then reversed, mapping all clusters to the microscope’s continuous coordinate space, in which clusters and cells can be co-located. The cluster map was finally validated by the introduction of a second set of hybridization probes, resulting in an overlay map with an accuracy of 2 microns (Fig. 1F).

**Figure 1.**
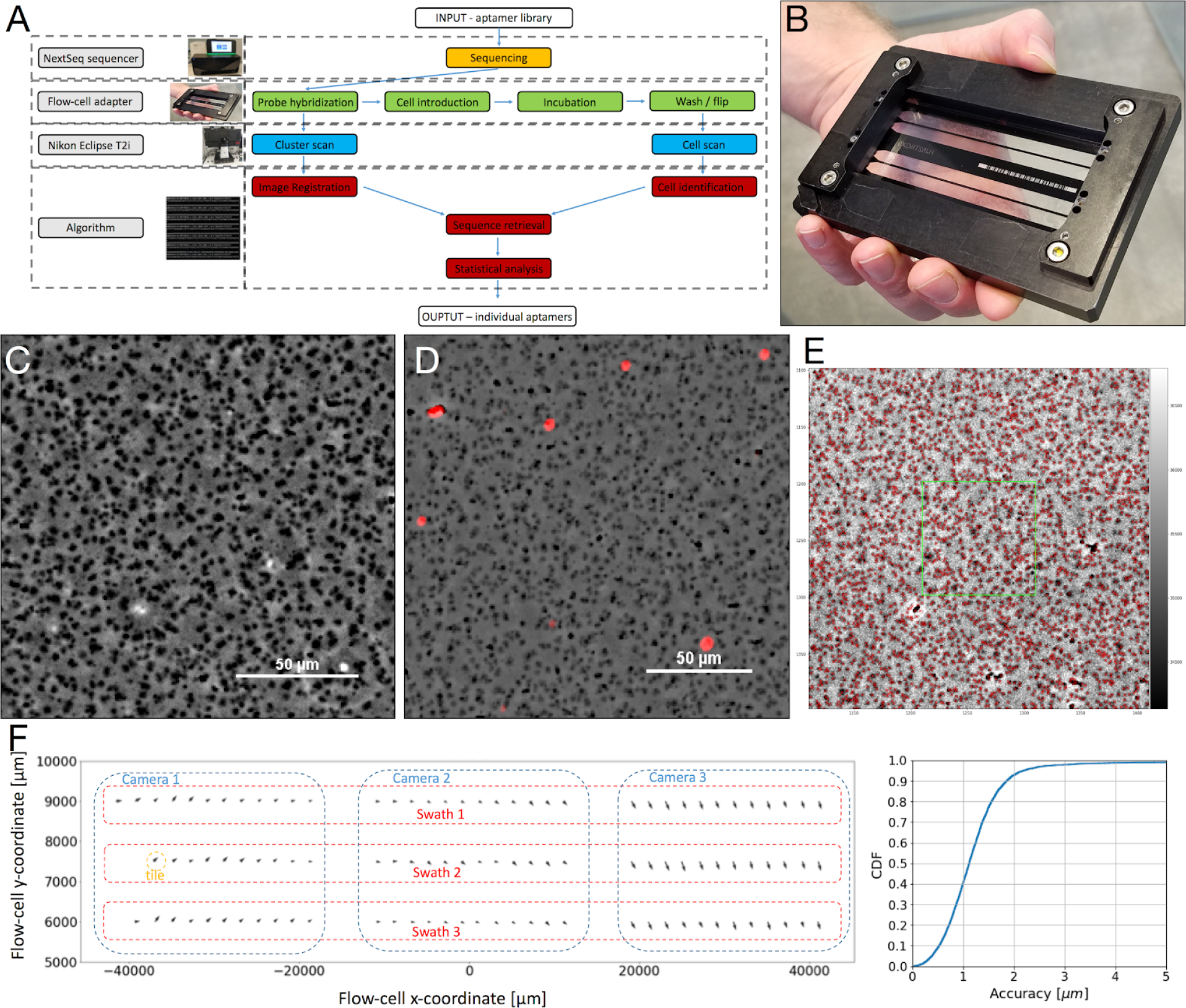
Imaging and registration of clusters in an external apparatus. A, a schematic representation of the platform. An input library is sequenced in a NextSeq 500 machine and the sequenced flow-cell proceeds to cellular assays and microscopic scan, concluding in an algorithm that highlights individual sequences. B, the mechanical adapter designed to support both microfluidics and microscopic imaging. C, a microscopic image of clusters captured in phase contrast microscopy, where the DNA clusters are visible in black (bar = 50 um). D, a microscopic image of fluorescently-labeled hybridization probes, overlayed on an image of clusters (bar = 50 um). E, a microscopic image of clusters captured in phase contrast microscopy, with red marks indicating where NextSeq clusters were mapped to, following the fine-tuned affine transformation. F, the accuracy of the reconstructed map. On the left, one lane with an arrow showing the average deviation vector per tile, based on fluorescent probes. On the right, the cumulative distribution function of the fluorescent probe deviation size. More than 90% of the probes are registered to less than 2μm of their actual location.

Once this process has been completed, the chip was washed with medium (see methods summary). Next, we introduced cap sequences into the chip. Cap sequences are oligonucleotides complementary to the constant stretches flanking the variable core of the library and are added in order to block the interference of those regions with the 3D structure of the folded oligonucleotide. Folding steps were done in medium with the presence of caps, with a folding protocol designed to match the selection environment of the specific original library. After the folding step, human cells were introduced into the chip (Fig. 2A). Various cell types were successfully introduced, with cells remaining viable for long periods of time (Fig. 2B). For cell recognition, we used a custom image processing workflow based on CellProfiler^34^, which yielded highly accurate recognition (Fig. 2C-E). At the end of this process, each cell could be successfully associated with a set of sequenced oligonucleotide clusters in the chip (Fig. 2F).

**Figure 2.**
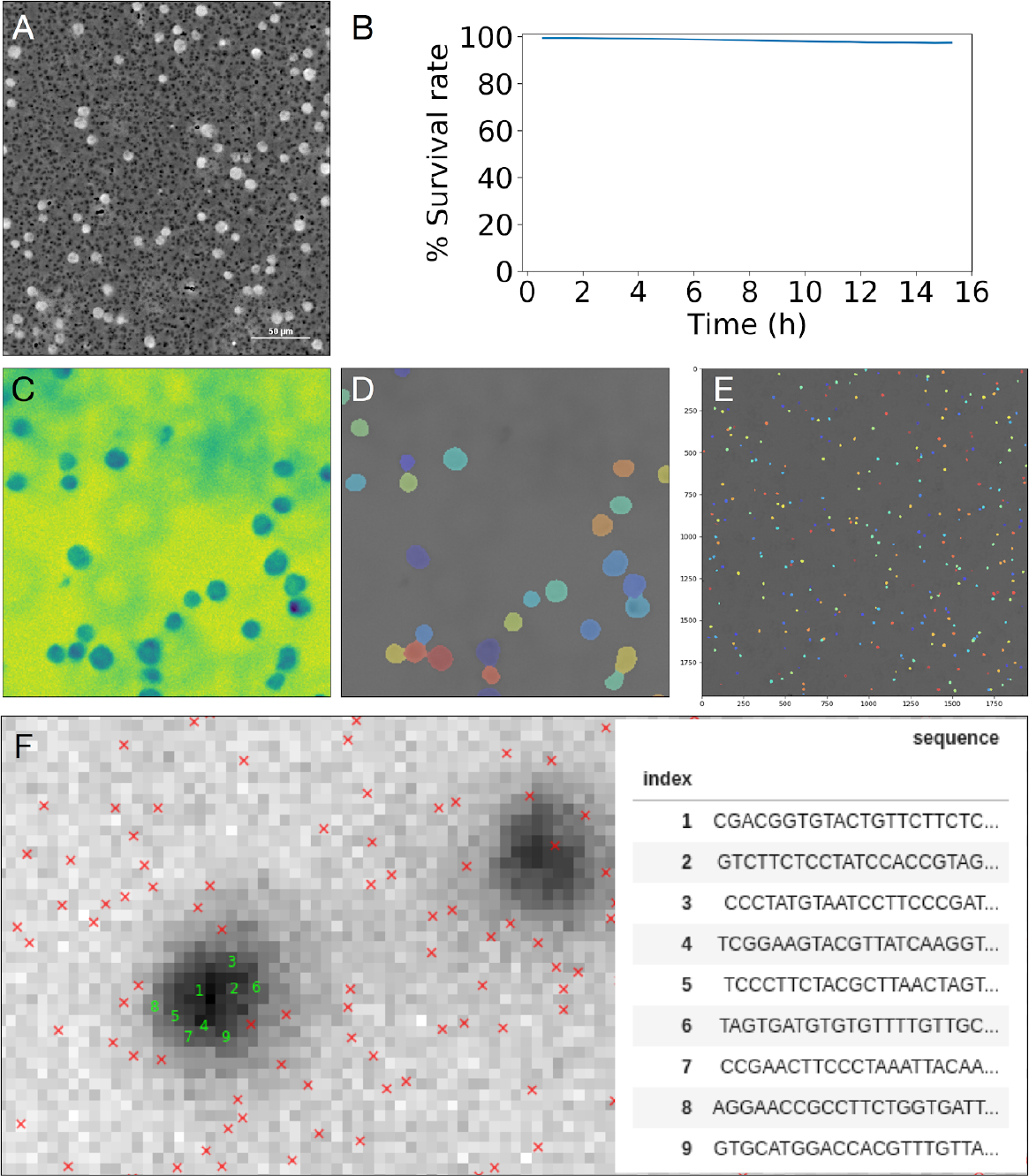
Target cell introduction and registration inside the flow-cell. A, a microscopic image of CCRF-CEM cells inside the flow-cell, overlayed on an image of clusters (bar = 50 um). B, survival over time of live MDA-MB-231 cells incubated inside the flow-cell, as indicated by Cas-3/7 staining. More than 95% of cells remain alive after 16 h. C, a partial field of CCRF-CEM cells as visualized by CellProfiler. D, the same area of C with cell identification by CellProfiler. E, a partial field with CellProfiler-identified, primary human AML cells. F. a highly magnified field containing two CCRF-CEM cells (dark areas) with marks for registered clusters in their vicinity. Nine of the clusters which co-localize with one of the cells are marked.

We first aimed to validate this platform by identifying known binding aptamers, using a chosen cell type, the human acute lymphoblastic leukemia (ALL) line CCRF-CEM. As a control, we planted into the sequencing process a previously-described aptamer to these cells, sgc8c^35^. Following sequencing, CCRF-CEM cells were introduced into the chip and incubated for up to 1.5 h in shaking. The chip was then flipped, causing unbound cells to physically fall out of focus while leaving only oligo-bound cells at the scanned surface (Fig. 3A, B). Above each bound cell (i.e. between the cell and the chip’s surface) are thus several clusters of candidate binding oligonucleotides. However, it is hard to determine directly which of these clusters - and therefore, which of the candidate oligonucleotides - is responsible for binding the cell. Since each binding candidate exists as multiple clusters throughout the chip, we counted the number of times one of its clusters co-localized with a bound cell. We then defined as ‘bound fraction’ the ratio of this number to the total number of clusters for that particular candidate (for example, a binding candidate would have a bound fraction of 0.2, if out of its total of 1,000 clusters, 200 were associated with bound cells). Bound fraction analysis confirmed a remarkable binding of CCRF-CEM cells by sgc8c (23-fold over background signal, *p* < 1.2×10^−4^) (Fig. 3C). Interestingly, when we repeated this experiment with sgc8c mutants planted into the sequencing process, the mutants exhibited a wide range of binding affinities manifested as bound-fraction values (Fig. 3D). To evaluate the reproducibility of these findings, we repeated this experiment comparing 4 different lanes from 2 different flow-cells. Results were highly reproducible and the correlation between lanes was 0.98 or higher (Fig. 3E).

**Figure 3.**
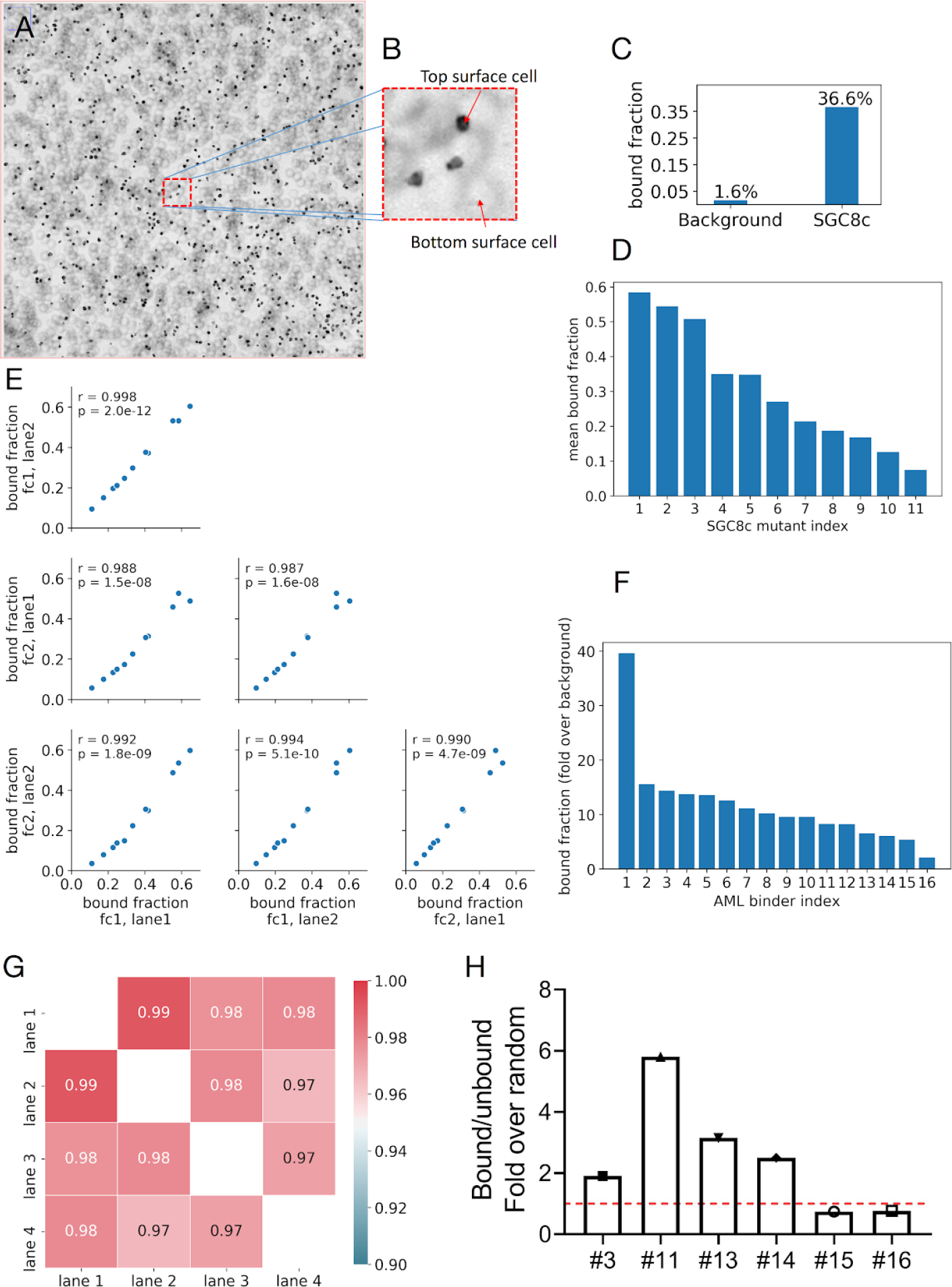
Analysis of cell-binding aptamers inside the flow-cell. A, CCRF-CEM cells in a flow-cell after incubation, wash, and flip. The top surface is in focus where the bound cells are visible in black, and the unbound cells laying on the bottom surface are visible as fuzzy rings. B, a magnified field from A. C, comparison of the bound fraction of SGC8c with the mean bound fraction across the entire flow-cell. SGC8c shows a 23-fold higher signal than the background. D, the top performing mutants of SGC8c, ranked by their bound fraction. E, correlation plots of SGC mutant bound fractions between 4 different lanes in 2 different flow-cells. The correlations demonstrate very high repeatability of the platform. F, binding aptamers to primary human AML cells, binding relative to background. G, correlation between 4 lanes in an experiment with primary human AML cells. H, performance of 4 leading primary human AML-binding aptamers, measured outside the flow-cell in an independent binding assay by quantitative PCR.

We next introduced acute myeloblastic leukemia (AML) cells, freshly isolated from a human donor, into a chip. Analysis revealed 16 aptamers with statistically significant bound fractions relative to background (*p* values ≤ 1.2×10^−4^) (Fig. 3F). Bound fractions were between 2 and 40 fold relative to background, and correlation of 0.965 or higher between lanes (Fig. 3G). The 4 leading aptamers performed well also in an independent binding assay (Fig. 3H).

We then proceeded to identify oligos that are directly cytotoxic towards target cells. This was also done on human triple negative breast cancer cells isolated from patient-derived xenograft mice^36^. As the reporter, we chose CellEvent caspase 3/7 green detection reagent^37^, a fluorogenic substrate of activated caspase 3/7 which signals apoptosis and was calibrated in the lab to work with these cells. In order to single out cells that went into the process already in poor conditions, we prestained the cells with Pacific Blue-labeled Annexin V. Immediately after staining, excess Annexin V was washed and thus labeling was restricted only to cells that went into the chip already in apoptosis. Time-lapse monitoring of these cells revealed the locations of cells undergoing apoptosis within as early as 3 hours (Fig. 4A, B), along with the locations of oligos inducing this effect.

**Figure 4.**
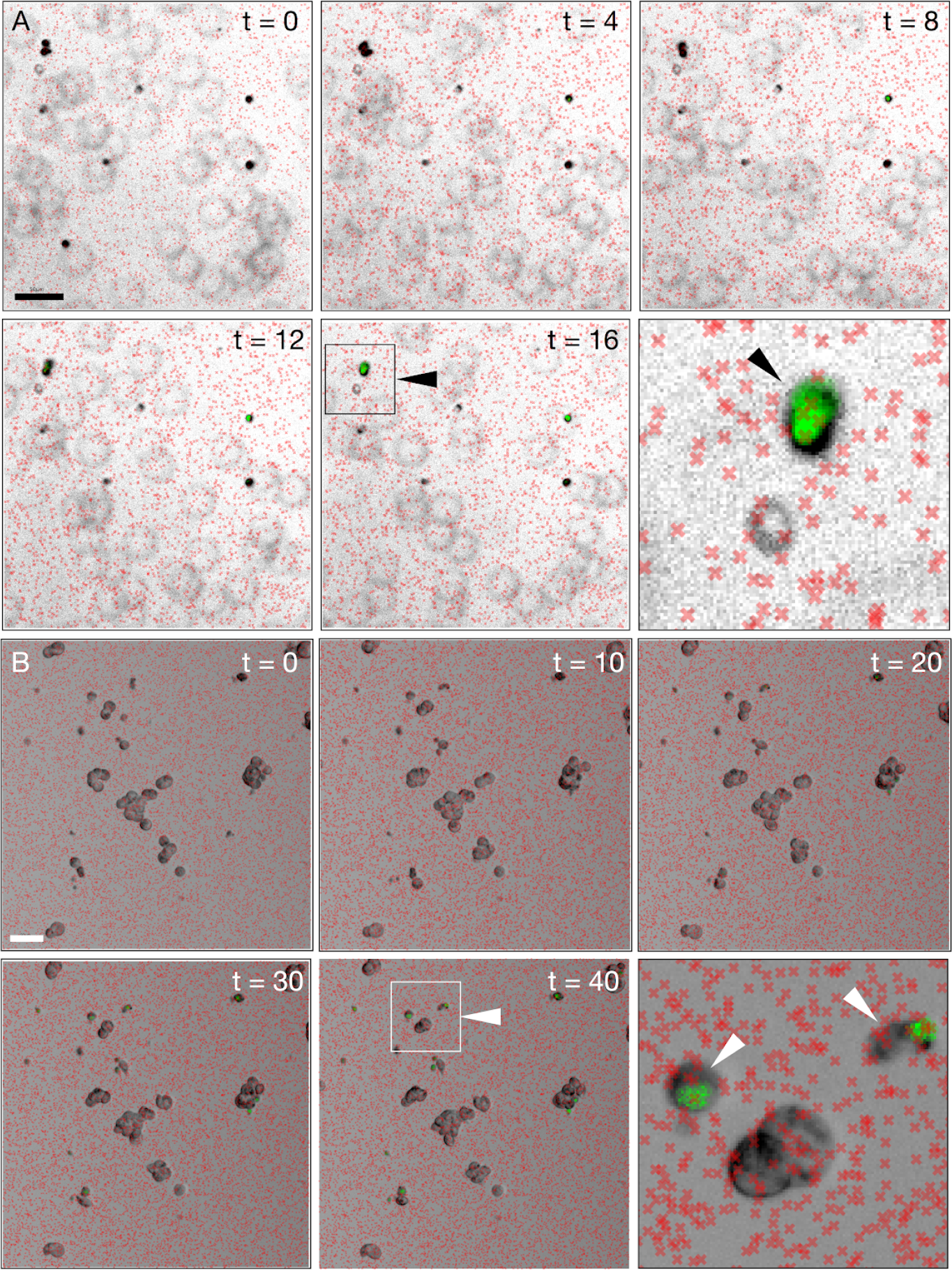
Discovery of tumoricidal oligonucleotides inside the flow-cell. AML cells and B, MDA-MB-231 cells, introduced into the flow-cell and incubated for the designated time periods (h). The last panel in each series shows a magnification of the framed region (square outline with arrowhead) from the previous panel. Arrowheads in the last panel indicate cells showing apoptosis, overlaid on the oligo clusters (red X marks) co-localized to them. Bars = 50 um.

## Discussion

In this paper, we draw a preliminary outline of a novel approach for drug discovery, in which therapeutic molecules - tumoricidal DNA oligonucleotides - are tailored *de-novo* for freshly acquired cell targets on a time scale of hours. This strategy incorporates several noteworthy, uncommon features for a therapeutic discovery technology. First, the process is driven by effect rather than by mechanism, and yields oligos that are directly tumoricidal against the target cells albeit through an unknown mechanism of action, although these oligos likely operate solely on the cell membrane and are not internalized, as this is built into the selection’s design. Experiments in our lab are underway to elucidate the span of potential mechanisms of action of the tumoricidal oligonucleotides, including on primary tumor cells with documented resistance to chemotherapy.

Second, the described strategy could incorporate selectivity into the selection process, by introducing ‘background’ or ‘non-target’ cells either into adjacent lanes or into same lanes as target cells but with discriminatory labeling. Statistical analysis would then score oligos based not only on the magnitude of their effect, but also on their target:off-target effect ratio, producing oligos that are not only effective but also potentially safe. Therapeutic safety should be determined much more rigorously than that, but installing this requirement early in the discovery process is highly favorable.

Third, every molecule resulting from this process is potentially unique. Given the immense variation within the human population, a per-target tailoring approach is arguably better than any form of generic treatment. It could also perform better than the ‘precision medicine’ (matching treatment based on patient-specific information or response) in the sense that it is not limited to a narrow set of drugs to select from. It is interesting to note that the tailored approach could (and, indeed, should) also include a ‘precision’ component,in which newly acquired targets are first tested against the entire set of previously-discovered oligos, and only if no hit is found that fits within specified performance margins, the target should proceed into the tailoring process. Introduction of this decision tree into the process could significantly enhance the platform’s effectiveness.

The process we describe here is carried out inside an external microscope. However, seamlessly performing it entirely inside the Illumina sequencer, following specific reconfiguration, is feasible, and would greatly improve the process and eliminate some of the problems associated with switching to an independent apparatus. In this regard, a sequencing machine could be converted into a miniaturized but powerful drug discovery machine.

Several challenges remain to be addressed. Most importantly, it is clear that conventional models of drug synthesis, testing, release, and regulation, are shaped for generic treatments, and do not support a tailored approach that emphasizes patient heterogeneity, rapid discovery, and per-target low cost. However, recently validated concepts could facilitate the necessary shifts. For example, on the technical level, therapies such as CAR-T cells, adoptive T cell transfers^38^, and dendritic cell vaccines, all fulfill the above-mentioned criteria and are based on fast, personalized processes. In addition, innovative regulatory models such as the *n*=1 clinical trial, are paving the ground for statistical analysis and approval of personalized treatments, and for specialized protocols for determining their safety rapidly and per patient. Once supported by the proper personalized regulatory models, the technology described here could arguably lead to a novel paradigm of personalized cancer therapy.

## Methods summary

Cell lines were purchased from American Type Cell Culture. Human primary acute myeloblastic leukemia (AML) cells were isolated from donors by standard procedures (IRB approval numbers 0297-15-TLV, 4573-17-SMC). Human (PDX-derived) triple negative breast cancer cells were a kind gift from B. Dekel, Sheba Medical Center, Israel. Target blasts were isolated by magnetic sorting using a commercial kit (Miltenyi Biotec) according to the manufacturer’s instructions. DNA libraries were designed with a 50-nt random core flanked by 20-nt constant regions and ordered from Integrated DNA Technologies. Randomization was done by hand mixing. All libraries passed in-house QC by HPLC before usage. Libraries were sequenced in an Illumina NextSeq 500 instrument with bleach replaced with HT1 buffer. Hybridization probes were designed to be 20-nt long and were sequenced together with the library. Probes were set to occupy ~0.1% of the flow cell clusters. Flow-cell scans were performed in a Nikon Eclipse Ti2-E microscope with Andor Zyla 4.2 camera, MRH20101 x10 objective, and Nikon Elements 5.0 software. Cluster imaging was done using phase-contrast microscopy. Cell imaging was done using Diascopic microscopy. Probe/markers imaging was done using fluorescent microscopy with a Chroma-49004 or Chroma-49006 filter cubes and Lumencor Sola SE II 365 illumination. Aptamer folding was carried out in folding buffer (PBS, 0.0045% w/v glucose, 1 mM MgCl2) for cell lines and LTC medium with 10% human serum for primary AML cells. Folding was done in the presence of 1 uM caps (complementary sequences to the constant regions, e.g. sequencing adapters and primers) by heating the chip to 95 °C for 10 min, cooling by incubation at 0 °C for 10 min, and another 10 min at 37 °C. 4,000,000 cells were divided into each lane and incubated for 1 h at 37 °C. Each of the candidates predicted by the flow cell assay was synthesized separately and tested in two different concentrations, 25 pM and 2.5 pM. After incubation, cells were washed in order to dilute the unbound oligos by 6 orders of magnitude. The bound fraction was then eluted from the cells and measured by qPCR.

## Statistical methods summary

The *p*-value was calculated by counting the number of oligonucleotide clusters whose center is within 3.0 um from the center of a cell and divided this by the total number of clusters. This value, the ‘mean bound fraction’, represents the fraction of clusters which are collocated with a cell. Next, for each DNA sequence, its ‘abundance’ was counted – how many oligonucleotide clusters on the flow-cell have this sequence, and its ‘bound cluster count’ – how many of its clusters are collocated with a cell. The raw *p*-value for a sequence is the binomial distribution survival with *p* being the mean-bound-fraction, *n* the sequence abundance, and *k* the bound-cluster-count. We ignored DNA sequences whose abundance was too low (< 5 oligonucleotide clusters) and counted the number of remaining DNA sequences. This number, the ‘sequence-count’, was used to correct for multiple experiments by a Sidak correction. Specifically, the corrected *p*-value for a sequence is defined to be 1 – (1 – *raw_p_value*)^*sequence_count*^.

## Acknowledgments

The authors wish to thank the team at Aummune Ltd (Sourasky Medical Center, Tel Aviv, Israel) for valuable technical assistance; to Prof. Benjamin Dekel (Sheba Medical Center) for the kind gift of TNBC cells.

## Author contribution

The following authors designed experiments, performed experiments, analyzed data and wrote the manuscript: NM, IR, GH, IB. The following authors designed experiments, performed experiments, and analyzed data: ZS, YA, EL, AZ, NB, NJ, MM. IB oversaw the project.

## Competing financial interests

The authors declare competing financial interests as follows. All authors are either employees and/or shareholders at Augmanity, a for-profit research organization that develops the technologies described in this article. The following authors are listed as inventors on patent applications related to technologies described in this article: IB, NM, IR, GH (PCT/IB18/00418; PCT/IB18/00613; US 62/738,235, pending).

